# Mora: abundance aware metagenomic read re-assignment for disentangling similar strains

**DOI:** 10.1101/2022.10.18.512733

**Authors:** Andrew Zheng, Jim Shaw, Yun William Yu

## Abstract

**Background:** Taxonomic classification of reads obtained by metagenomic sequencing is often a first step for understanding a microbial community, but correctly assigning sequencing reads to the strain or sub-species level has remained a challenging computational problem.

**Results:** We introduce Mora, a MetagenOmic read Re-Assignment algorithm capable of assigning short and long metagenomic reads with high precision, even at the strain level. Mora is able to accurately re-assign reads by first estimating abundances through an expectation-maximization algorithm and then utilizing abundance information to re-assign query reads. The key idea behind Mora is to maximize read re-assignment qualities *while simultaneously* minimizing the difference from estimated abundance levels, allowing Mora to avoid over assigning reads to the same genomes. On simulated diverse reads, this allows Mora to achieve F1 scores comparable to other algorithms while having less runtime. However, Mora significantly outshines other algorithms on very similar reads. We show that the high penalty of over assigning reads to a common reference genome allows Mora to accurately infer correct strains for real data in the form of short E. coli reads and long Covid-19 reads.

**Conclusions:** Mora is a fast and accurate read re-assignment algorithm that is modularized, allowing it to be incorporated into general metagenomics and genomics workflows. It is freely available at https://github.com/AfZheng126/MORA.

## 1 Background

When analyzing microbial communities through metagenomic sequencing, a fundamental task is to determine which reference genome a specific sequencing read originates from [1]. This gives information about microbial composition and allows for mapping-based analysis of genetic variation. A common first step in a processing pipeline is to use a fast taxonomic read classifier such as Kraken2 [2], CLARK [3], Centrifuge [4] or others. While such methods are extremely fast, they are not sensitive enough to assign reads to the strain level. Since strain-level resolution has important functional implications [5–7], sensitive methods that are able to resolve reads at the strain level are needed.

To resolve reads at the level of strains, a naive approach would be to more sensitively align the read to a set of candidate reference genomes using a read aligner such as Bowtie2 [8], Minimap2 [9], or CORA [10] (no relation to Mora) and then take the best reference genome as the correct assignment. However, strain-level reference genomes share large regions of similarity, so many ambiguous mappings are inevitable. To overcome this limitation, one can statistically calculate abundance information of the candidate set of reference strains and only use references that have high enough abundances [11–14]. Afterwards, one can re-assign reads to the “correct” references, i.e. reference strains that seem to be abundant in the metagenomic sample.

However, re-assigning reads to the correct references while maintaining abundance estimates is non-trivial. For example, if one were to assign a multi-mapped read to the most abundant reference strain with a putative mapping, all reads coming from a region of similarity between two strains will be mapped to only one strain—in this case the most abundant one. This is not an accurate assignment of the reads and will skew the abundance. Importantly, not all algorithms do abundance estimation *and* read assignment; some algorithms calculate only abundances and do not output re-assigned reads [15–17].

In this paper, we present Mora, a tool that allows for sensitive yet efficient metagenomic read re-assignment and abundance calculation at the strain level for both long and short reads. Given an alignment in SAM or BAM format and a set of reference strains, Mora calculates the abundance of each reference strain present in the sample and re-assigns the reads to the correct reference strain in a way such that abundance estimates are preserved. We rigorously formulate this problem as an optimization problem and give provable guarantees on our heuristic algorithm. We show that Mora is more effective than Pathoscope2 [12], a state-of-the-art read re-assigner, at disambiguating similar strains present in a sample on simulated data for short reads while being an order of magnitude faster. Furthermore, we show that Mora has similar if not better F1 scores compared to Pathoscope2, Kraken2, and Clark while taking less time and RAM on simulated long read data. We then verify our results on real long-read data as well.

## 2 Results and discussion

### 2.1 Pipeline of Mora

Mora’s pipeline consists of two main steps (Figure 1): abundance estimation and read re-assignment. Like other metagenomic abundance estimators [17, 18], it first utilizes a standard generative probabilistic model on input mappings and performs inference using the expectation maximization (EM) algorithm, augmented with a set cover algorithm to filter out spuriously abundant genomes [17]. Mora’s novelty comes from the subsequent step, where we re-assign reads in a manner *dependent on the calculated abundances*. Mora models the problem of maximizing correct read assignments while minimizing the difference between predicted abundance levels and final abundance levels as a non-linear minimization problem (Section 3.3 and Figure 2). Although we use a greedy heuristic, we prove that heuristic is guaranteed to improve the minimization score. Finally, Mora outputs a (re-)assignment of each read to a reference genome that is consistent with the calculated abundances.

**Fig. 1:**
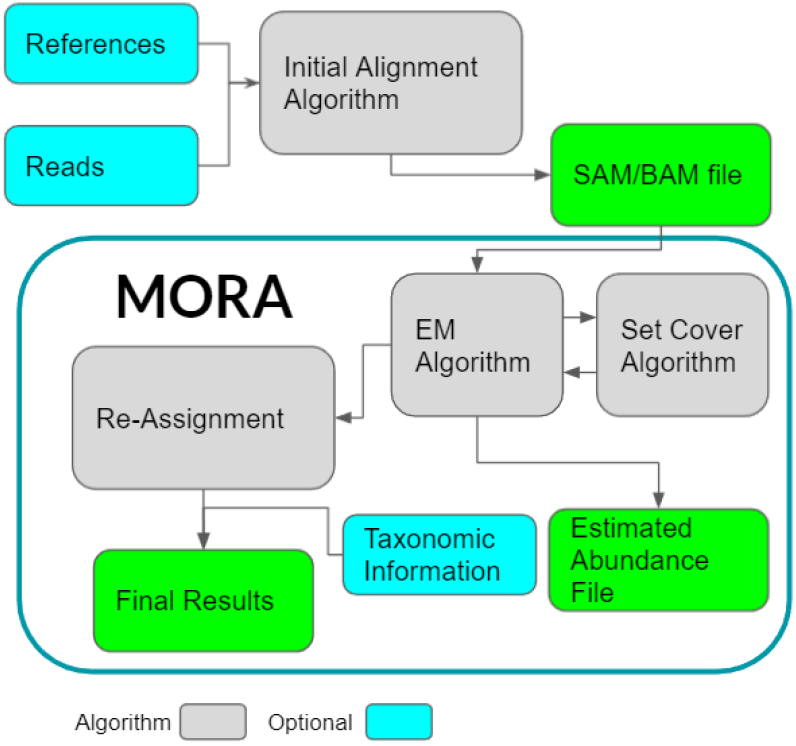
The pipeline for Mora to output re-assignments from query and reference FASTA files. Mora’s processing steps are enclosed by the dark green rectangle. The abundance estimation step includes the Expectation Maximization (EM) algorithm and a set cover algorithm to filter out spuriously abundant genomes [17]. The reassignment step uses the estimated abundance from the previous step to re-assign reads.

**Fig. 2:**
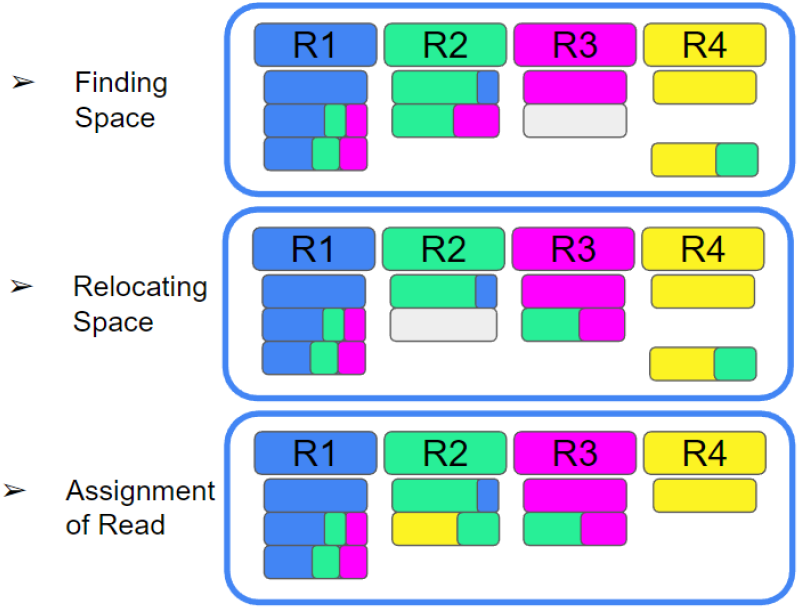
An example of Mora’s approach to read re-assignment. The exact algorithm is outlined in Section 3.3. R1-R4 are four reference genomes labelled with colours, and the 8 reads shown have colour content proportional to the mapping scores with respect to each coloured reference genome. Grey boxes are not reads, but available read assignments based on Mora’s estimated abundances. Step 1: Assigning reads based on highest mapping scores leads to invalid assignments, since R4 can only store 1 read based on the abundance constraint. Step 2: we move a more ambiguously assigned read in R2 to R3 instead, opening space in R2. Step 3: We move the multi-assigned read in R4 to R2, where there is now space.

### 2.2 Benchmarking details

We benchmarked Mora using simulated short and long reads against Pathoscope2 [12], Kraken2 [2], and Clark [3]. Pathoscope2 can identify strains, compute abundance levels, perform re-assignments, and output informative result summaries. It has seen usage in numerous studies [19–22], and its algorithm has also been incorporated into other taxonomic classification pipelines [23]. Kraken2 and Clark are two taxonomic classifiers that classify reads by assigning them a taxonomic ID. In the case of Kraken2, it may classify reads to the least common ancestor of the possible taxons to reduce the chance of false assignments. For Clark, the taxonomic IDs can be manipulated for strain-level assignments. This was also attempted for Kraken2, but errors were encountered when trying to build a custom database. Hence, Kraken2 was unable to work at the strain level. All algorithms except Pathoscope2 were run using their default parameters, which would not run due to the high number of strains. Instead, certain parts in Pathoscope2’s code had to be augmented to allow it run correctly. Its initial library construction code no longer works and must be replaced with code from MetaScope [24]. The output step also had to be fixed to allow for large number of reads. These changes did not affect how Pathoscope2 re-assigns reads, so the only change should be being able to run on large datasets and a decrease in runtime. We now denote the augmented Pathoscope2 as AugPatho2, and the augmented code can be found in same repository as the appendix files. Given these caveats, Pathoscope2 appears to be the current practical state-of-the-art.

We also considered MetaMaps [14] and Sigma [13], two other re-assignment tools, but they were not included in this comparison as the software are no longer actively maintained and parts of both software are no longer functional; other studies also had issues with these two programs [15, 25].

F1 score, sensitivity, and precision are used to evaluate the accuracy of the final read re-assignment at three different taxonomic ranks: strain, species, and genus (see section 3.4). For the purpose of this paper, two DNA sequences are of the same species/genus if their NCBI taxID corresponds to the same species/genus name. Two DNA sequences are of the same strain if their accession number is the same.

### 2.3 Re-Assignment of reads to similar strains

To test strain identification, 58 E. coli reference genomes were obtained from NCBI and 950,000 short 150bp pair-end reads were generated for 3 of those strains, so each strain has about 30x coverage. We aligned the reads with Pufferfish/Puffaligner [26], a very efficient read alignment method. We then used Mora and AugPatho2 to reassign the reads, and we report the number of assigned reads for each strain. We also performed assignment using Clark and Bowtie2 with their default parameters. The results are shown in Figure 3. AugPatho2 is programmed to perform an initial assignment by calling Bowtie2 with pathoMAP. To allow the use of other alignment algorithms, we generated the SAM file independently of AugPatho2 and ran pathoID and pathoMAP on it. Since all the strains had the same taxonomic ID, Kraken2 could not be benchmarked on this dataset. To run Clark, the taxonomic IDs were changed to be unique from each other.

**Fig. 3:**
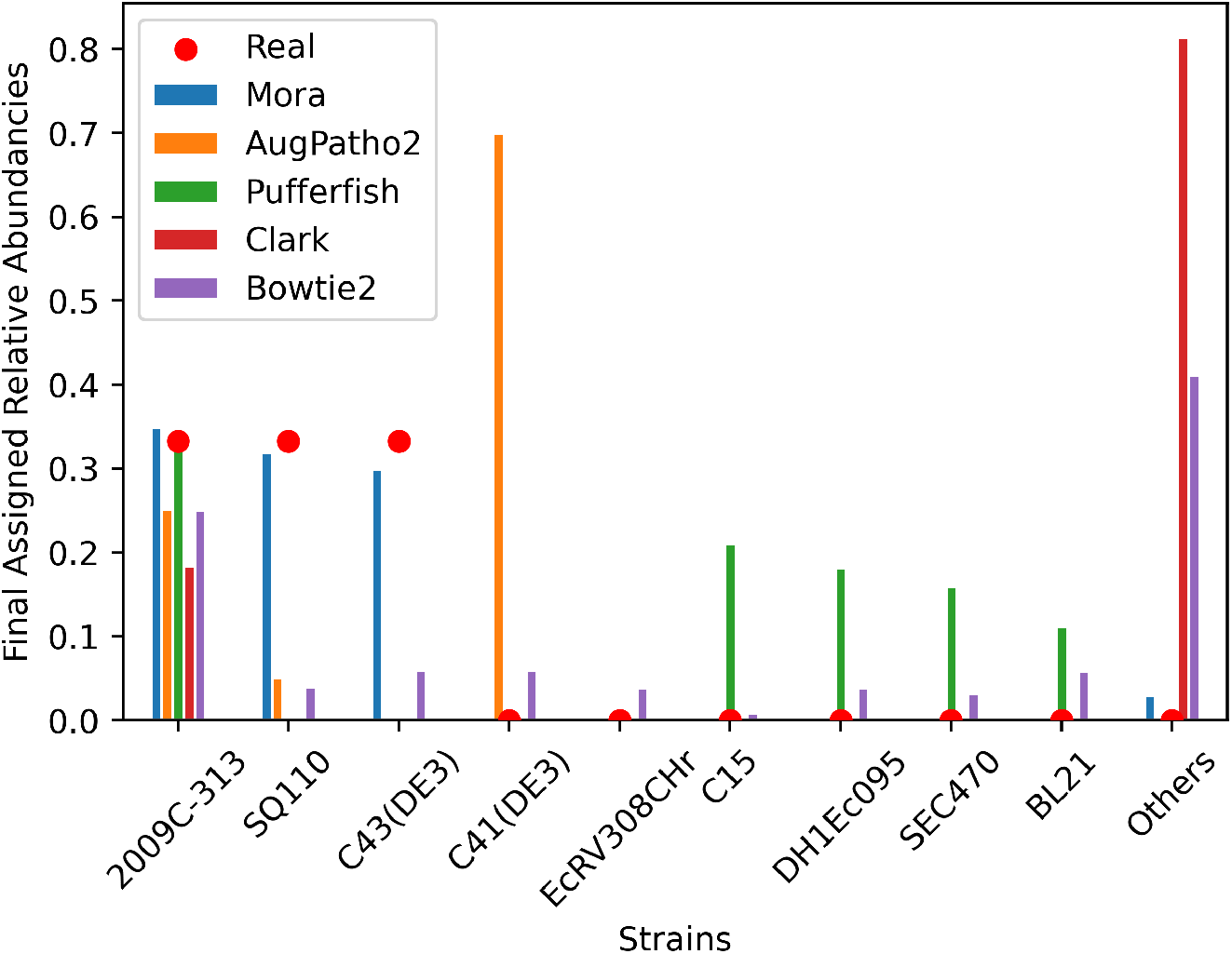
Relative assignment abundancies of Mora, AugPatho2, and Pufferfish of 950,000 synthetic short 150bp pair-end reads to 58 E. coli strains. The synthetic short reads were simulated from the three E. coli strains: 2009C-3133, SQ110, and C43(DE3). The strains listed had at least 2000 assigned reads from at least one of the algorithms. Relative assignment percentages of the final assignments from three different algorithms are represented by the different coloured bars. The real abundance levels of the strains are represented by the red dots. AugPatho2 is Pathoscope2 but with slight changes in the code to make it able to run and output results. Assignment by Pufferfish and Bowtie2 were done by choosing the primary alignment in the SAM file, while assignment by Mora and AugPatho2 was done using Pufferfish as the initial aligner. Clark and Bowtie2 were run without any additional algorithms. Mora’s assignments are closest to the real abundancies.

In Figure 3, Mora, AugPatho2, and Pufferfish were able to identify the presence of all three strains. Mora had a F1 score of 74.03, much higher than 29.01, 32.48, 18.35, 32.28, the F1 scores of AugPatho2, Pufferfish, Clark, and Bowtie2 respectively. AugPatho2 mapped most reads to the strain C41(DE3), a strain whose average nucleotide identity to C43(DE3) is 99.94% according to OrthoANI [27]. This may be caused by how Pathoscope2’s algorithm incorporates reference length into its calculations of read alignment scores. The length of strain C43(DE3) was 56,061 base pairs shorter than strain C41(DE3), making the alignment scores to C41(DE3) higher than those to C43(DE3), which is the likely reason for a large number false assignments. As most high mapping scores were equal in value, selecting the primary alignment without regard of abundance levels is likely the reason for Pufferfish’s high false positive rate. Clark was unable to align 81% of the reads while Bowtie2 assigned almost equal amounts of reads to all strains. The number of reads assigned to each strain can be found in appendix table A.1.

### 2.4 Large-scale simulated short read re-assignment at the species and genus level

We used the complex Illumina 400 dataset of 400 different microbial genomes and its corresponding simulated 26.6 million 75bp paired-end short reads [28] to test the accuracy of short read alignment. We consider two cases: when there is a good guess of which genomes the reads come from, and when there is no information at all. For the first case, we built a reference library (REF-1) from the Illumina 400 dataset (400 genomes) that was used to simulate our reads. For the second case, we built a reference library (REF-2) using the complete bacterial genomes from NCBI RefSeq (6487 genomes). Importantly, REF-2 does not contain all references in REF-1. REF-2 contains 36% of the strains, 72% of the species, and 87% of the genera of REF-1. Due to the lack of species and genus level taxonomic data for REF-1, the scores for species and genus are worse than the scores for strain. When using REF-2 as the reference genomes, scores for the strain rank are not reported due to the low number of strains from REF-1 included in REF-2.

When aligning to REF-1, Pufferfish was unable to map 13.7 million reads while Bowtie2 was unable to map 5.7 million. When aligning to REF-2, Pufferfish was unable to map 15.8 million reads while Bowtie2 was unable to map 10.1 million.

As shown in Tables tables 3 and 4, AugPatho2 performed worse than Mora when aligning to REF-2 using Bowtie2, but was slightly better in other cases. However, it was much slower in runtime due to the pathoREP module in AugPatho2. For large data sets, the pathoREP module wasn’t able to output the final XML file due to lack of memory. We had to split the SAM file into 3-5 smaller files and change the XML output script to accommodate this problem. As seen in Table 1, Mora was over 19 times faster than AugPatho2, while only having a 3 times increase in RAM usage. Using Bowtie2 increased scores significantly due to its ability to map more reads than Pufferfish, however, this did lead to a large increase in computer runtime. Clark had comparable but slightly better scores than Mora when using Bowtie2 in all categories except on REF-2 at the genus level. Kraken2 on the other hand performed worse than Mora when using Bowtie2 on REF-1 but had slightly better scores on REF-2 at the genus level.

**Table 1:** Wall clock time (s) and maximum RAM usage (GB) for algorithms on aligning simulated short reads (SR)

| Algorithm | SR to REF-1 |  | SR to REF-2 |  |
| --- | --- | --- | --- | --- |
|  | Time(s) | RAM(GB) | Time(s) | RAM(GB) |
| Mora | 381.48 | 47.04 | 514.30 | 46.16 |
| AugPatho2 | 28103.34 | 51.58 | 28111.39 | 15.99 |
| Pufferfish | 89.70 | 6.791 | 273.22 | 8.388 |
| Bowtie2 | 490.55 | 1.591 | 753.09 | 21.265 |
| Kraken2 | 73.48 | 2.40 | 70.54 | 24.33 |
| Clark | 222.25 | 41.09 | 311.63 | 91.10 |

Other than using more RAM on short reads to REF-2, Mora consistently used fewer resources than AugPatho2 and was substantially faster. Time cannot be directly compared between the Mora/AugPatho2 and Kraken/Clark as they require the output from one of the initial aligners. CPU time and other information can be found in appendix table A.4. All algorithms were run with 8 threads on a Ubuntu 20.04.4 LTS server with an AMD Ryzen 3970X 32-core CPU with 256 GB of RAM.

### 2.5 Re-Assignment of long reads

To test Mora on long reads, we used badreads [29] v0.4.0 with default parameters on REF-1 to simulate 1.37 million long nanopore reads. When using Minimap2 to perform the initial alignment to REF-1, 44.3 thousand reads could not be aligned. When using Minimap2 to align to REF-2, 183 thousand reads could not be aligned.

Tables tables 5 and 6 lists the scores evaluating assignment accuracy at the strain, species, and genus level of long reads for the different algorithms. Mora consistently had higher scores on long reads for REF-1, though this lead was lost when looking at the scores for REF-2. Though AugPatho2 had a very similar score to Mora, Table 2 shows a large difference in memory and RAM usage. Mora performs at least four times faster and uses up to 27.45 fold less RAM compared to AugPatho2 on long reads. The main bottleneck for AugPatho2 was again PathoREP, the algorithm to write its result files. Even after the augmentations of PathoREP to reduce its runtime and memory usage, AugPatho2 still had higher runtime and RAM usage than Mora for long reads. Both Kraken2 and Clark had lower F1 scores than Mora in both REF-1 and REF-2, which is expected as they were initially designed for short reads. Furthermore, Kraken2’s tendency to assign common ancestry when dealing with multi-mapping reads results in very low scores at the species level.

**Table 2:**
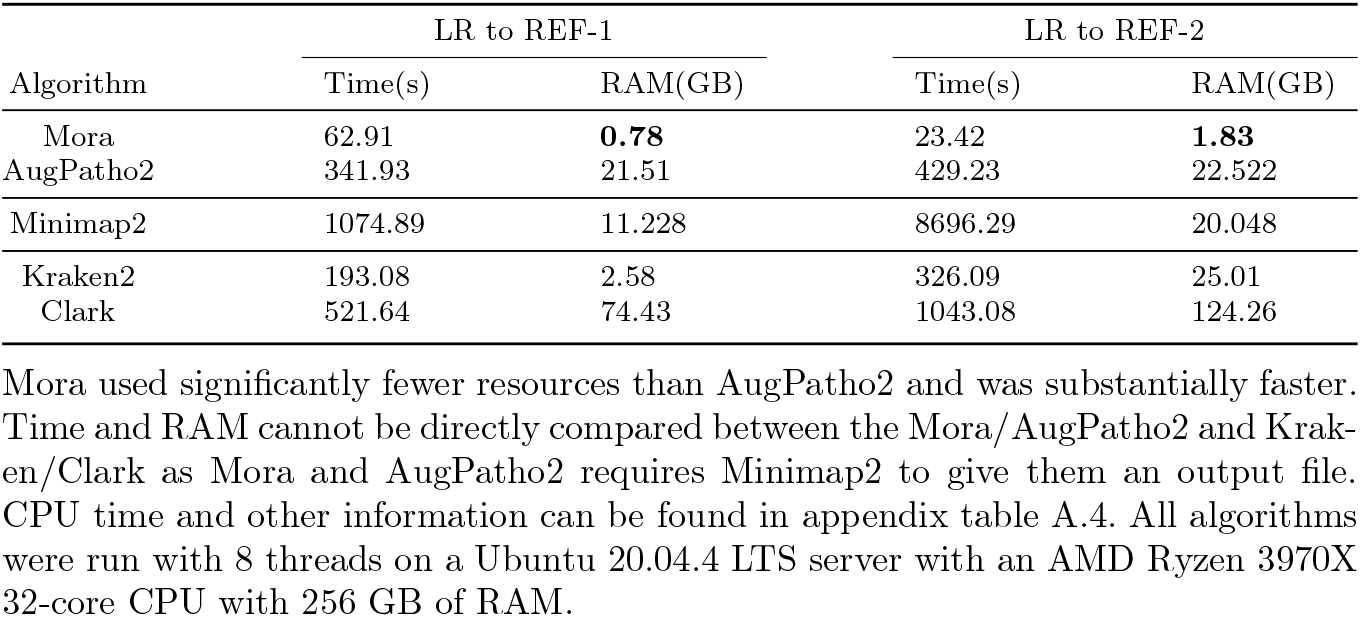
Wall clock time (s) and maximum RAM usage (GB) for algorithms on aligning simulated long reads (LR)

**Table 3:** Scores of algorithms when aligning short reads to REF-1

| Algorithm | Strain |  |  | Species |  |  | Genus |  |  |
| --- | --- | --- | --- | --- | --- | --- | --- | --- | --- |
|  | F1 | Sensitivity | Precision | F1 | Sensitivity | Precision | F1 | Sensitivity | Precision |
| Pufferfish | 60.58 | 44.05 | 96.94 | 59.17 | 42.53 | 97.19 | 65.00 | 48.19 | 99.80 |
| Mora + Pufferfish | 63.51 | 47.21 | 97.02 | 59.22 | 42.56 | 97.27 | 65.00 | 48.21 | 99.83 |
| AugPatho2 + Pufferfish | 63.55 | 47.24 | 97.07 | 59.23 | 42.57 | 97.29 | 65.02 | 48.21 | 99.83 |
| Bowtie2 | 83.13 | 74.29 | 94.36 | 78.31 | 66.87 | 94.46 | 87.25 | 77.71 | 99.46 |
| Mora + Bowtie2 | 85.41 | 76.33 | 96.95 | 80.56 | 68.79 | 97.18 | 87.53 | 77.96 | 99.77 |
| AugPatho2 + Bowtie2 | **90.21** | **84.22** | 97.12 | **85.33** | **75.93** | 97.39 | 92.31 | **85.87** | **99.79** |
| Kraken2 | NA | NA | NA | 0.29 | 0.27 | 0.33 | 87.17 | 79.26 | 96.84 |
| Clark | 86.88 | 78.10 | **97.87** | 82.82 | 71.46 | **98.47** | **97.85** | 78.68 | 99.46 |

Both Mora and AugPatho2 were able to improve the scores of the initial aligner, though AugPatho2 appears to have a slight systematic edge.

**Table 4:** Scores of algorithms when aligning short reads to REF-2

| Algorithm | Species |  |  | Genus |  |  |
| --- | --- | --- | --- | --- | --- | --- |
|  | F1 | Sensitivity | Precision | F1 | Sensitivity | Precision |
| Pufferfish | 26.32 | 16.56 | **64.14** | 40.32 | 25.55 | 92.08 |
| Mora + Pufferfish | 34.73 | 24.10 | 62.15 | 54.44 | 38.26 | 94.34 |
| AugPatho2 + Pufferfish | 35.59 | 24.70 | 63.69 | 54.97 | 38.62 | 95.26 |
| Bowtie2 | 44.51 | 35.55 | 59.52 | 71.83 | 58.22 | 93.73 |
| Mora + Bowtie2 | 46.54 | 37.17 | 62.23 | 72.42 | 58.71 | **94.51** |
| AugPatho2 + Bowtie2 | 39.14 | 28.79 | 61.11 | 62.72 | 46.98 | 94.28 |
| Kraken2 | 14.95 | 13.03 | 17.53 | **78.51** | **68.42** | 92.08 |
| Clark | **49.89** | **40.94** | 63.85 | 69.65 | 59.10 | 86.94 |

**Table 5:**
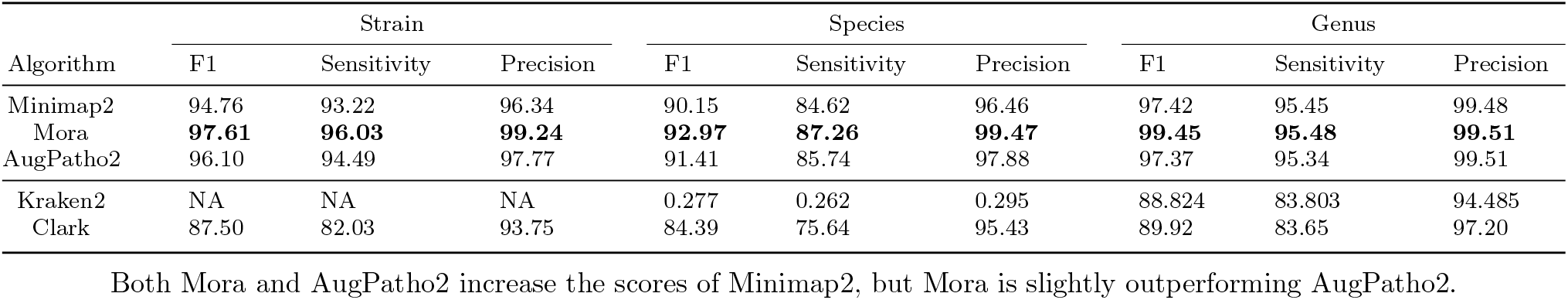
Scores between Minimap2, Mora, AugPatho2, Kraken2, and Clark when aligning long simulated reads to REF-1

| Algorithm | Strain |  |  | Species |  |  | Genus |  |  |
| --- | --- | --- | --- | --- | --- | --- | --- | --- | --- |
|  | F1 | Sensitivity | Precision | F1 | Sensitivity | Precision | F1 | Sensitivity | Precision |
| Minimap2 | 94.76 | 93.22 | 96.34 | 90.15 | 84.62 | 96.46 | 97.42 | 95.45 | 99.48 |
| Mora | **97.61** | **96.03** | **99.24** | **92.97** | **87.26** | **99.47** | **99.45** | **95.48** | **99.51** |
| AugPatho2 | 96.10 | 94.49 | 97.77 | 91.41 | 85.74 | 97.88 | 97.37 | 95.34 | 99.51 |
| Kraken2 | NA | NA | NA | 0.277 | 0.262 | 0.295 | 88.824 | 83.803 | 94.485 |
| Clark | 87.50 | 82.03 | 93.75 | 84.39 | 75.64 | 95.43 | 89.92 | 83.65 | 97.20 |

Both Mora and AugPatho2 increase the scores of Minimap2, but Mora is slightly outperforming AugPatho2.

**Table 6:** Scores between Minimap2, Mora, AugPatho2, Kraken2, and Clark when aligning long simulated reads to REF-2

| Algorithm | Species |  |  | Genus |  |  |
| --- | --- | --- | --- | --- | --- | --- |
|  | F1 | Sensitivity | Precision | F1 | Sensitivity | Precision |
| Minimap2 | 52.02 | 46.60 | 58.86 | 83.60 | 77.74 | 90.42 |
| Mora | 53.84 | 48.23 | 60.92 | 84.64 | 70.70 | 91.55 |
| AugPatho2 | **54.30** | **48.31** | **61.97** | **85.40** | **78.78** | **93.23** |
| Kraken2 | 16.45 | 15.39 | 17.66 | 79.56 | 74.43 | 85.47 |
| Clark | 49.78 | 43.17 | 58.78 | 73.37 | 65.66 | 83.12 |

Both Mora and AugPatho2 increase the scores of Minimap2, but AugPatho2 is slightly outperforming Mora.

### 2.6 Results on real sequencing data

To show applicability to real sequencing data, we ran Mora on two sets of real data, each representing a different scenario.

In the first case, we look at real short E. coli reads from three different assemblies. Mora and AugPatho2 (with Bowtie2 as the initial aligner) were run on 30,000 pair-end short reads of average length 250 bp. These 30,000 reads were composed of 10,000 reads from three SRA runs representing the assembly genomes of INF13/18/A, INF191/17/A, and INF32/16/A (see Appendix table A.7), while the references were the 58 E. coli strains used previously with the addition of the three new strains. The average nucleotide identity (ANI), calculated by skani [30], of the three strains to each other were between 96.4 and 96.57 (see Appendix table A.7). As seen in Figure 4, Mora was able to assign most reads to the three INF strains, while AugPatho2 assigned a lot of reads to a different complete genome assembly: JJ2434. The ANI between JJ2434 and INF191/17/A was 99.39, but its complete genome (i.e. one single contig) sequence length was 3.4 million bps longer than the longest scaffold in the assembly for INF191/17/A. This is the likely reason for the large error in AugPatho2, which uses reference lengths for scoring. Similar to other experiments, Bowtie2 assigns a decent amount of reads to all E. coli strains. Mora had an F1 score of 62.34 while AugPatho2 and Bowtie2 had F1 scores of 49.48 and 41.10 respectively.

**Fig. 4:**
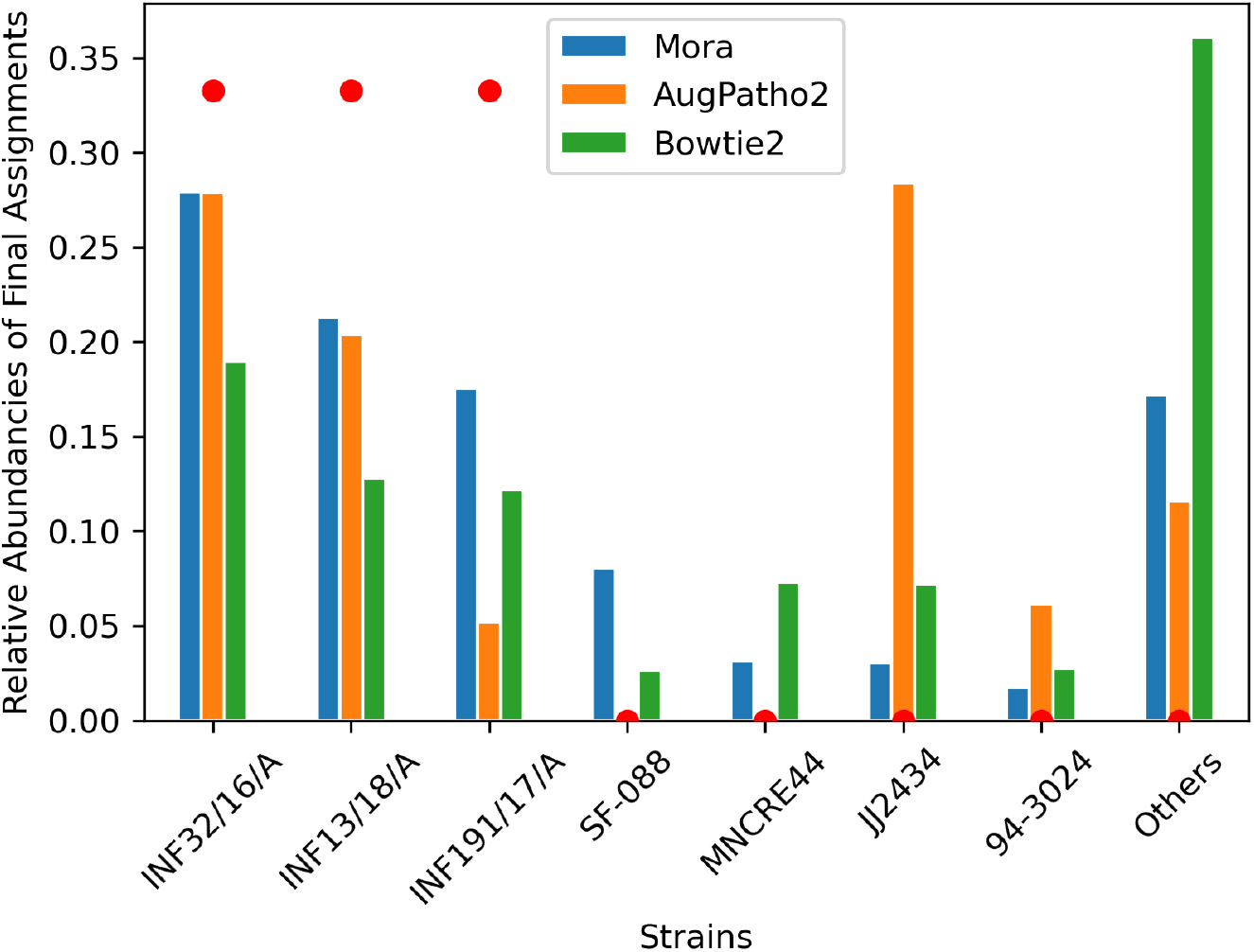
Comparison of assignment percentages of real short E. coli reads from the three assemblies: INF32/16/A, INF13/18/A, and INF191/17/A by Mora, AugPatho2, and the initial aligner Bowtie2. The real abundance levels of the strains are represented by the red dots. Mora assigns a lot more reads to the INF assemblies compared to AugPatho2 and Bowtie2.

In the second case, we still test Mora on very similar strains, but now the reads are long PacBio Covid-19 reads. Mora and AugPatho2 (with Minimap2 as the initial aligner) were used to re-assign 2635 Covid-19 reads of average length 1239 bp to five different Covid-19 strains: Wuhan, Alpha, Beta, Delta, and Omnicron. The specific sequence used to represent these strains was chosen by taking the first sequence for the Pango lineage as shown on NCBI Virus. For the Alpha strain, two different sequences (one from USA and one from Germany) were selected, but their ANI was 99.91, meaning they differed by around 30 nucleotides. ANI between the different strains was between 99.50 and 99.85. For more sample details, see Appendix table A.8. As the samples were taken from USA during 2021, which was when the Alpha strain was the most prevalent, it is likely that the samples are likely to belong the Alpha strain. As seen in Figure 5, Mora assigns the most reads to Alpha, with AugPatho2 assigning the second most number of reads to Alpha. Based on our initial knowledge of the samples, we conclude that the results from Mora are more likely to be correct in this experiment.

**Fig. 5:**
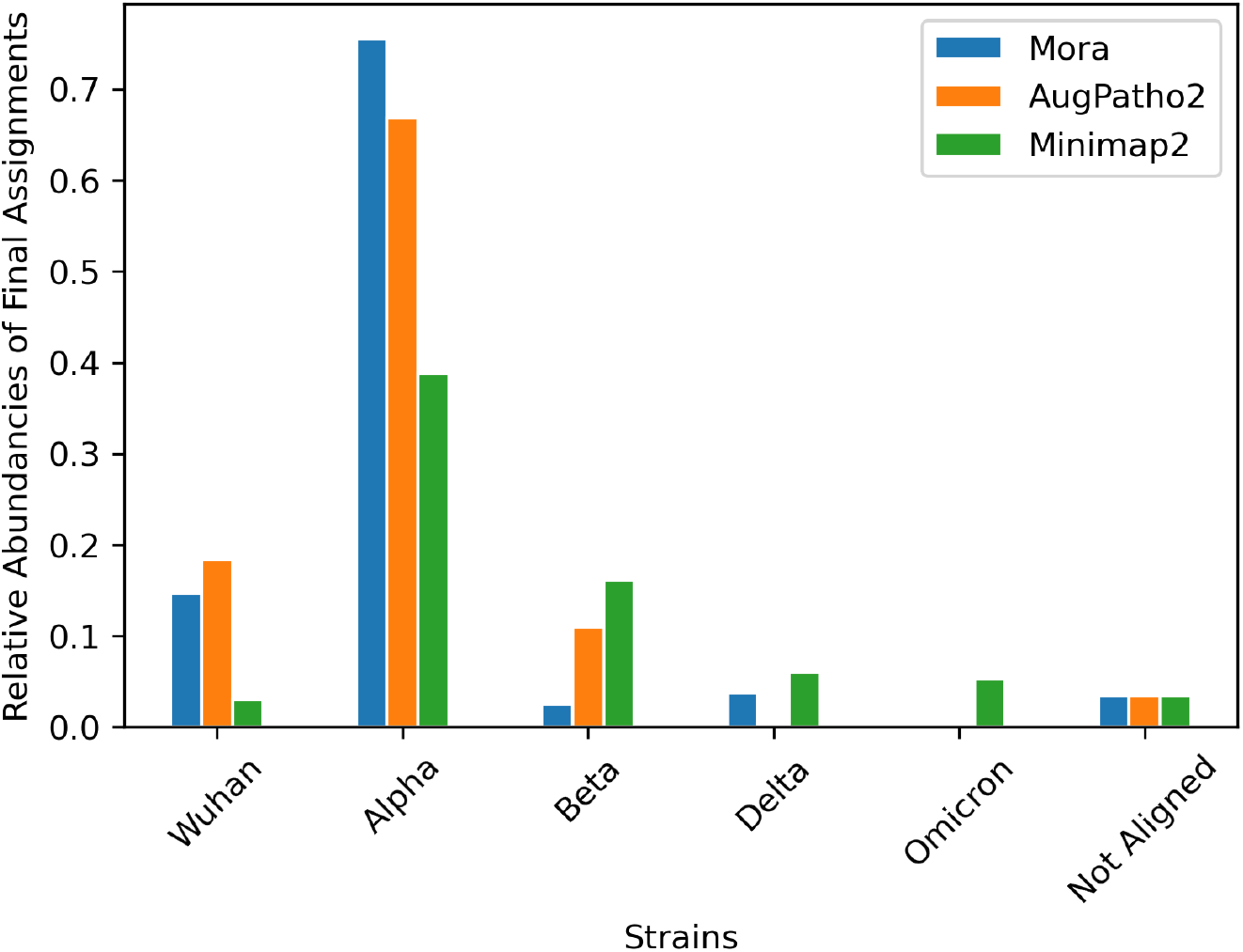
Final abundances of Mora, AugPatho2, and Minimap2 of covid-19 samples. Long PacBio Covid-19 reads were from 48 individuals from Kentucky, USA published in June 2021. All algorithms assign most of the reads to the Alpha strain, which based on time and location is the most likely strain, with Mora assigning the most.

## 3 Methods

We define *mapping score* as some positive value associated to each alignment that measures how good the alignment is. In practice, we use the AS:i secondary flag that is present in most read aligners, but our theory holds for any such score.

### 3.1 Initial read mapping

The input to Mora is a SAM/BAM file with an initial alignment of the reads. After filtering out unaligned reads, the remaining reads are fed into Mora. At minimum, Mora only requires a SAM file with header information to output read (re)assignments. Mora optionally allows for the inclusion of taxonomic information of reference genomes in the final output by using accession and taxonomic information from NCBI.

For convenience, Mora is also implemented with a read mapping module using Snakemake [31], allowing the user to use Mora from a set of input reads and references. The mapping modules allow the user to use Pufferfish, Bowtie2, or Minimap2. However other programs can be used as long as their generated SAM contains a header with the reference genomes and the AS:i secondary mapping flag is available in each alignment record. After the SAM file is generated, the reads that could not be assigned are filtered out. These reads are later added to the final output with the “NO ALIGNMENT” string being assigned to them.

### 3.2 Abundance estimation

We solve the problem of estimating the abundance levels of reference genomes by adapting the common model from calculating RNA transcript abundances used by Agamemnon [17] and RSEM [32]. In this model, each read is generated by first selecting a reference and then a position on that reference. Assuming the same model for all the reads, we have the following likelihood function for observing a set of reads from a set of reference genomes.

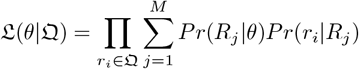

where M is the number of references, 𝔔 is the set of reads, *θ* is the abundance estimation, *Pr*(*R*_*j*_|*θ*) is the prior probability of selecting reference *R*_*j*_, and *Pr*(*r*_*i*_|*R*_*j*_) is the conditional probability of generating read *r*_*i*_ from reference *R*_*j*_.

*Pr*(*r*_*i*_|*R*_*j*_) is computed by normalizing the mapping score between read *r*_*i*_ and reference *R*_*j*_ over all mapping scores from *r*_*i*_. The estimation of *θ* is done with an EM algorithm. To reduce the number of iterations needed by the EM algorithm to converge, the reads are reduced to a set of equivalent classes where two reads 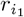 and 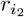 are equivalent if they align to the same set of references.

After every 10 iterations, we implement Agamemnon’s idea of using a set cover algorithm to remove redundant references with low abundances from the rest of the iteration process. This results in better-estimated abundances of leftover references while also increasing the convergence rate. These algorithms are implemented based on the Cedar algorithm used by Agamemnon. For more information, please consult Agamemnon’s original paper.

### 3.3 Re-Assignment of reads

Mora then adds to the functions of Agamemnon by using the estimated abundance levels to perform read re-assignment. The assignment of reads based on their mapping scores while trying to stay true to the estimated abundance levels can be modeled as a variant of the Weapon-Target Assignment (WTA) problem [33]. Given different weapons that are to be fired on a set of different targets, the objective is to find which weapons should be assigned to which targets to minimize the expected remaining health of the targets. Formulating it as a non-linear problem, let {*T*_1_, *T*_2_, …, *T*_*N*_ } be the set of targets and {*W*_1_, *W*_2_, …, *W*_*M*_ } be the set of different weapons. For each weapon type *W*_*i*_, there are *w*_*i*_ number of it and each has probability *P*_*ij*_ to destroy the target *T*_*j*_. After assigning *X*_*ij*_ of weapon *W*_*i*_ to target *T*_*j*_, the probability that *T*_*j*_ survives is 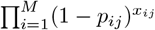. Thus the WTA problem aims to minimize the following non-linear problem with constraints:

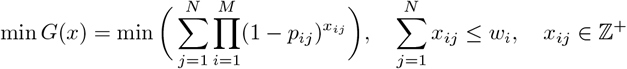

Instead of selecting which weapons to assign to which targets to maximize expected damage, we are selecting which reads to assign to which references to maximize likelihood. Let *R*_*j*_ represent a reference genome, *r*_*i*_ represent a read, and *P*_*ij*_ represent the probability that *r*_*i*_ comes from *R*_*j*_ is true, calculated in the same way as for abundance estimation. To model our metagenomic problem as the WTA problem, we have three assumptions:

1. Every reference *R*_*j*_ appears *a*_*j*_ · *M* times in the set of references, where *a*_*j*_ is the estimated abundance of *R*_*j*_ and *M* is the total number of reads. The value of *a*_*j*_ · *M* is approximated to the nearest integer.
2. Every read maps to some reference genome.
3. Every read is unique, so *w*_*i*_ = 1 for *i* = 1, …, *M*.

These three assumptions result in the re-assignment problem being formulated as the following minimization problem with constraints:

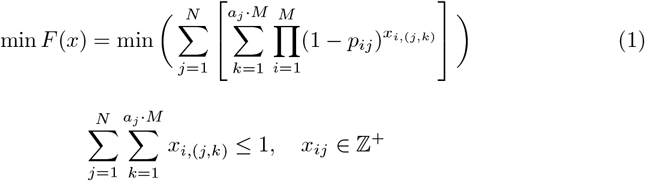

The second sum shows that there are *a*_*j*_ · *M* of reference *M*_*j*_ and the *xi*,(*j,k*) is how much of read *r*_*i*_ we assign to the *k*th copy of reference *M*_*j*_. This model is more punishing against undershooting compared to overshooting as over-assigning reads to a reference does not decrease the objective function *F*(*x*). This is desirable as it is better to identify all the low-abundance genomes and undershoot the most abundant genome than to not identify the low-abundance genomes. As leaving a read un-assigned and assigning it to a very wrong reference both contribute a value of 1 to the objective function *F*(*x*), the assumption that every read maps to some reference is needed to prevent large numbers of false positives. Using Equation 1, we can now use optimization methods to get exact solutions, though this is not very practical given the bad runtime scaling for large data sets.

As this is an NP-hard problem [33], Mora uses a greedy algorithm that finds a relatively good solution. Mora views each reference as a bin with a fixed space capacity. Every time a read is assigned to a reference, the available capacity of the reference decreases. Once the capacity of a reference is full, no other read can be assigned to that reference unless something is taken out. By default, the capacity *C*_*j*_ of a reference *R*_*j*_ is *C*_*j*_ = *a*_*j*_ + 0.001. The amount of space each read takes up is 1/*M*. As reads get assigned to the references, the references *R*_*i*_ can be represented as a list of assignments 𝒜(*R*_*i*_) containing the reads that have already been assigned to it.

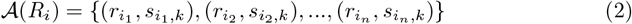

where 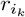 represents a read, 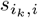 is the corresponding mapping score between 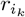 and *R*_*i*_, and *n* is the current number of reads that have been assigned to *R*_*i*_. The list is ordered using 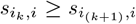 and the capacity limitation is 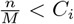. If 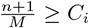 we say that the reference *R*_*i*_ is full. Similarly, the reads *r*_*j*_ can be represented as a list of potential mappings ℳ(*r*_*j*_).

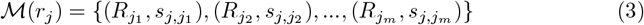

where the list is ordered such that 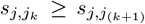. The total score of a read *r*_*j*_ is defined to be

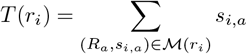

which gives us, by definition, that 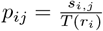.

Mora assigns the reads in terms of priority. A read *r*_*i*_ is given priority 1 if the read maps to only one reference. A read *r*_*i*_ is given priority 2 if the ratio of the second best score to the best score is less than a threshold. By default, this threshold is 0.5 but can be changed. A read is given priority 3 if it doesn’t satisfy the conditions of being priority 1 or priority 2. Once priority values are assigned, the priority 1 reads are assigned, followed by priority 2 reads, and then priority 3 reads.

Priority 1 reads are assigned to the unique reference they map to. For priority 2 reads, they are first sorted from highest best mapping score to lowest best mapping scores. In this order, the reads are assigned to the reference with the best mapping score if that reference has space. If the reference is at full capacity, the read is relabeled as a priority 3 read. When assigning priority 3 reads, all mappings between the priority 3 reads and references that still have space are sorted in terms of the score into a list. The reads are then assigned in order of this list, or left over for a second round of assignment if all of its potential references are full. After the initial assignment is done, Mora will try to “open up space” in a reference to assign leftover reads.

#### Definition 1.

*For a read r*_*i*_ *and* 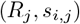 ∈ ℳ(*r*_*i*_), *R*_*j*_ *can open up space for r*_*i*_ *if it is full and there exists a* 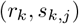 ∈ 𝒜(*R*_*j*_) *such that r*_*k*_ *can be moved to another reference* 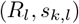 ∈ ℳ(*r*_*k*_) *with the condition that*

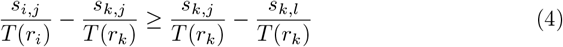

Using the notation in this definition, we have the following theorem.

#### Theorem 1.

*If r*_*i*_ *is a read and R*_*j*_ *is a reference that can open up space for r*_*i*_ *by re-assigning r*_*k*_ *to another reference R*_*l*_, *then doing so and then assigning r*_*i*_ *to R*_*j*_ *decreases the value of F* (*x*) *from Equation 1*.

*Proof*. The act of performing this re-assigning *r*_*k*_ from *R*_*j*_ to *R*_*l*_ and then assigning *r*_*i*_ to *R*_*j*_ is equivalent to changing from [*x*_*ij*_, *x*_*kj*_, *x*_*kl*_] = [0, 1, 0] to 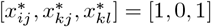. Since 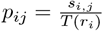, the last condition of being able to open up space gives us that

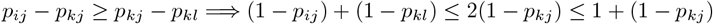

The right side can be written as

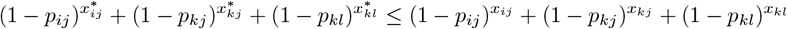

Since there is space in *R*_*l*_, changing the three values *x*_*ij*_, *x*_*kj*_, *x*_*kl*_ doesn’t result in the invalidation of any constraints and doesn’t affect any other terms in the sum of *F*(*x*). Thus, performing the re-assignment causes a decrease in the value of *F*(*x*). □ □

If space cannot be opened up in *R*_*j*1_, Mora will try to open up space in *R*_*j*2_, and so on. If space cannot be opened for any of the references the read maps to, the read will be left to the end to be assigned randomly with weights corresponding to the mapping scores. At this stage, mapping these reads using probability is plausible as their mapping scores are relatively similar to each other. A simple example of this greedy algorithm is shown in Figure 2.

### 3.4 Evaluation metrics

Genomes are classified as the same species or genus depending on the taxonomic information listed in NCBI. Taxonomic information of the data is obtained from NCBI Taxonomy’s FTP database (/pub/taxonomy/accession2taxid/) using the live and dead nucleotide sequence records. Genomes are classified as the same strain if their accession numbers are the same. The calculation of the accuracy metrics only considers reads that were successfully aligned by the first assignment algorithm. This allows us to evaluate the re-assignment algorithms without having the results be affected by the first assignment algorithms.

Read assignment accuracy on the simulated data sets is measured using F1 score, sensitivity, and precision. Let *r*_*i*_ be a read generated from a reference *R*_*i*_. At any taxonomic rank, *r*_*i*_ is labeled as a true positive if it is mapped to reference *R*_*j*_ that agrees with *R*_*i*_ at that taxonomic rank. If *R*_*i*_ and *R*_*j*_ do not agree at that rank, then *r*_*i*_ is labeled a false positive. If *r*_*i*_ is not assigned to anything, it is not labeled as anything. For example, a read generated from *Escherichia coli* with accession number CP0001, assigning it to *Escherichia coli* with accession number CP0005 would be a true positive for the species and genus rank, but a false positive at the strain rank. Assigning it to *Escherichia fergusonii* with accession number CP1001 would be a true positive at the genus rank, but a false positive at the strain and species rank.

For a taxonomic rank, let TP be the total number of true positives for that rank and let FP be the total number of false positives for that rank. We define

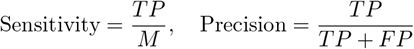

where *M* is the total number of reads. The F1 score is defined to be the harmonic mean of sensitivity and precision.

### 3.5 Data Simulation and Availability

30 E. coli genomes assemblies were downloaded from NCBI Assembly and combined to form 58 E. coli strains. The three strains 2009C-3133, SQ110, and C43(DE3) were chosen randomly to simulate short reads 1.37 million 150bp pair-end short reads. The reads were simulated according to a uniform distribution using art illumina [34] with the default parameters for pair-end reads. The simulated 26.6 million 75bp pair-end short read data from REF-1 was obtained from [28], where it simulated using iMESS Illumina with a skewed distribution. For the simulation of long reads, Badread [29] was used with the default parameters corresponding to mediocre Oxford nanopore reads with quantity 20x. The read distribution is proportional to the length of the references.

The real E.coli data for the strains INF32/16/A, INF191/17/A, and INF13/18/A can be found at SRR15443628, SRR15497613, and SRR10587526 respectively. The Covid-19 reference strains were taken from NCBI Virus by searching for their respective accession codes (found in Appendix table A.8) while the samples were taken from SRR14752036.

For a use-case, the *E. coli* reads and references are available at https://github.com/AfZheng126/MORA-data. The full appendix tables of scores, time, and memory usage for the different simulations are also available in the same repository as the *E. coli* reads/reference data.

## 4 Conclusion

In this work, we presented Mora, a new flexible algorithm and pipeline for assigning reads at the strain level. Mora takes as input an alignment file and re-assigns the reads to strains by (1) estimating abundance information and (2) modelling the reassignment problem as a discrete non-linear minimization problem for which Mora’s heuristic solution has provable guarantees. We showed that Mora performs well compared to other read assignment algorithms, but truly shines on reads from very similar strains.

Additionally, we showed that Mora is fast and practical to use, even on large datasets, with speeds and memory usage several times better than AugPatho2. Though the speed of the full pipeline is slower than Kraken2 and Clark, it makes up for it this with higher F1 scores at the strain, species, and genus levels on long reads and very similar reads.

We found that there is a surprising lack of well-engineered tools that deal with the specific problem of sensitive read re-assignment to the strain level. Thus we designed Mora using general mathematical formulations, leading to it working well on many kinds of data. Furthermore, Mora is engineered to be modular and easy to use—the minimal input required is just a single SAM/BAM file. Thus we believe that Mora will be a useful tool for researchers interested in studying strain-level read information from metagenomic sequencing data.

### Supplementary information

Download the Appendix Tables.xlsx file from https://github.com/AfZheng126/MORA-data to view all supplementary tables. The code for AugPatho2 can also be found in the same repository.

## Declarations

### Availability of data and materials

All analysis software written for this manuscript are available in the https://github.com/AfZheng126/MORA repository, which contains the Mora software described in this manuscript. The software requires Rust ¿1.60.0 and has been tested on Linux environments. It is available under an MIT-style free and open source license.

The datasets supporting the conclusions of this article are available in the https://github.com/AfZheng126/MORA-data repository.

### Authors’ contributions

The study was conceived by J.S. and supervised by Y.W.Y. A.Z. wrote the software and performed the experiments.

### Consent for publication

This manuscript has been seen and approved by all listed authors.

### Ethics approval

Not applicable.

### Consent to participate

Not applicable.

